## Supplementary File 1120 for "Articular cartilage corefucosylation regulates tissue resilience in osteoarthritis"

**Supplemental Material**

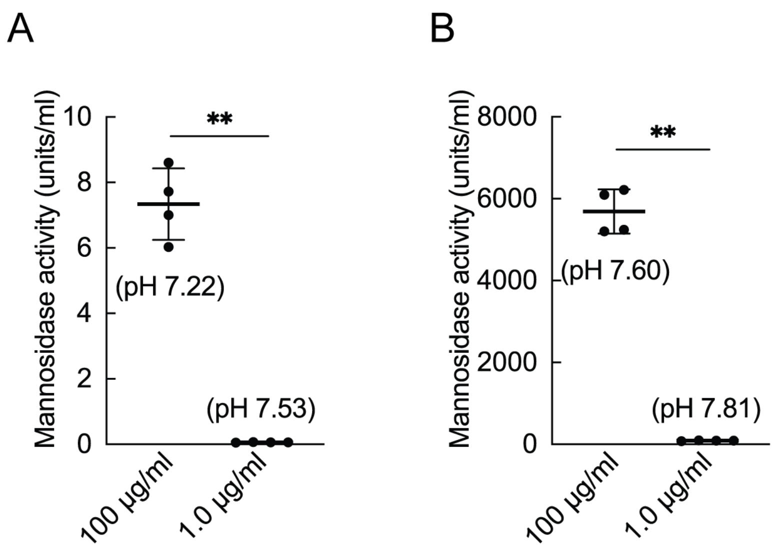


**S1 Figure. Enzymatic activity under physiologic pH conditions in joint injection and cartilage culture systems**

The activity of mannosidase under saline (A) and DMEM (B) condition at 37 ºC (n = 4). Data are shown as mean ± standard deviation. **P < 0.01. the Welch t-test was used for statistical analysis.

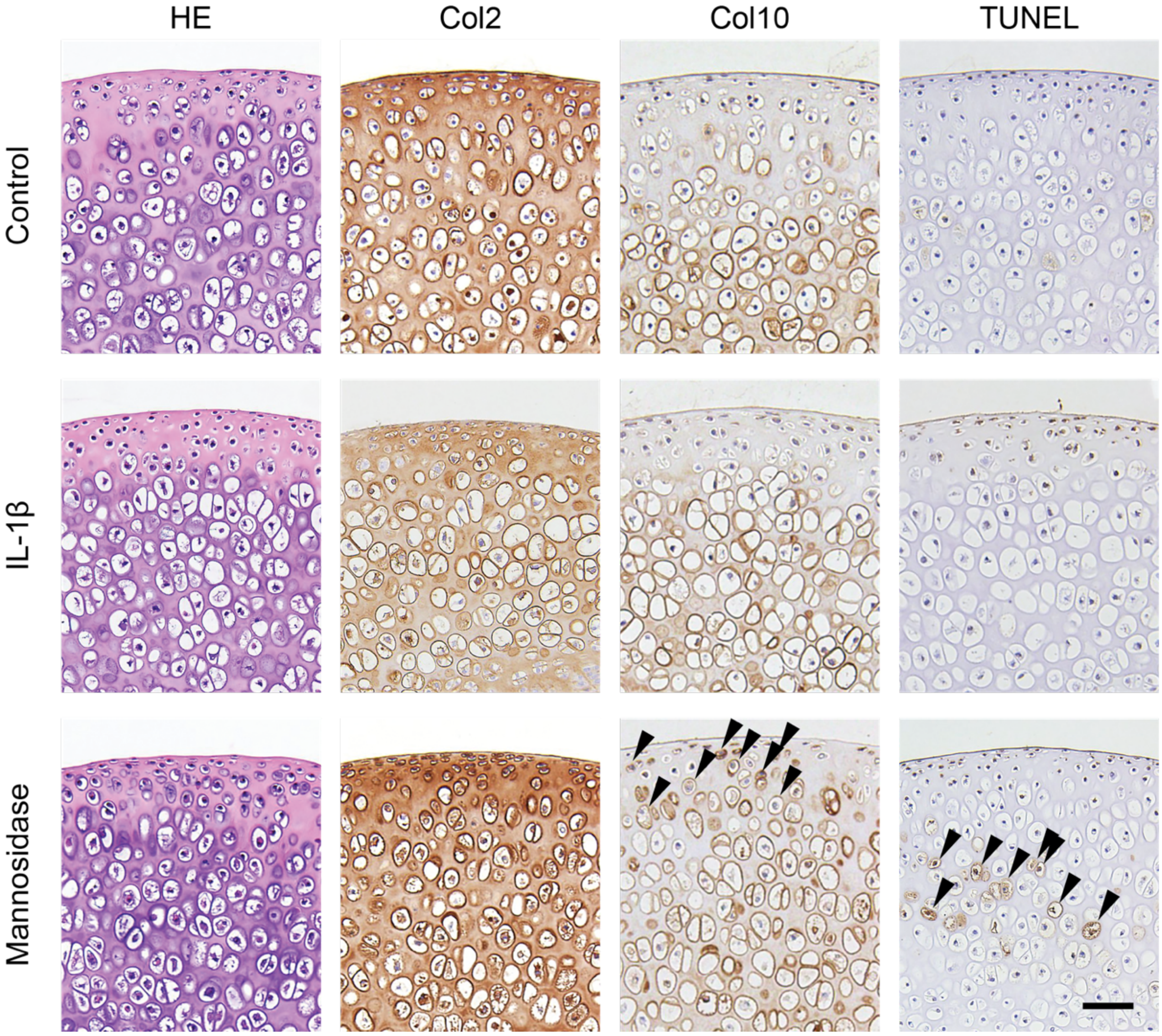


**S2 Figure. Histological effects of mannosidase on cartilage culture systems**

Mannosidase-treated cartilage was stained with HE, and immunohistochemical staining for Col2, Col10 and TUNEL. Overall, Col2 staining decreased in the IL-1β-treated cartilage. The superficial layer of cartilage affected by mannosidase contained Col10-positive cells, whereas the intermediate layer contained TUNEL-positive cells (black arrow). Scale bars, 50 μm. HE, hematoxylin and eosin; Col2, type II collagen; Col10, type X collagen; TUNEL, Terminal deoxynucleotidyl transferase dUTP nick end labeling; IL, interleukin.

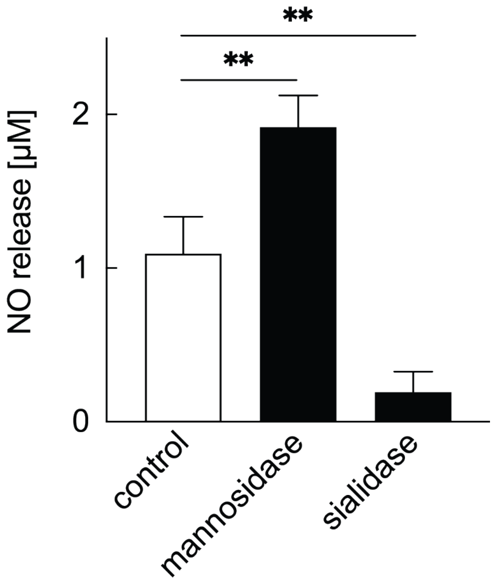


**S3 Figure. Impact of other glycoside hydrolases on NO production from cartilage**

NO release in articular cartilage explants. Explants were cultured under normal culture conditions (control) and under the effect of mannosidase (1.9 U/mL) and sialidase (1.9 U/mL). NO release was photometrically measured using Griess reagent in the culture supernatant. **P < 0.01 versus the control group. n = 4 samples (8 mice per group). One-way ANOVA with the Dunnett multiple-comparison test was used to perform statistical analysis. NO, nitric oxide; ANOVA, analysis of variance.

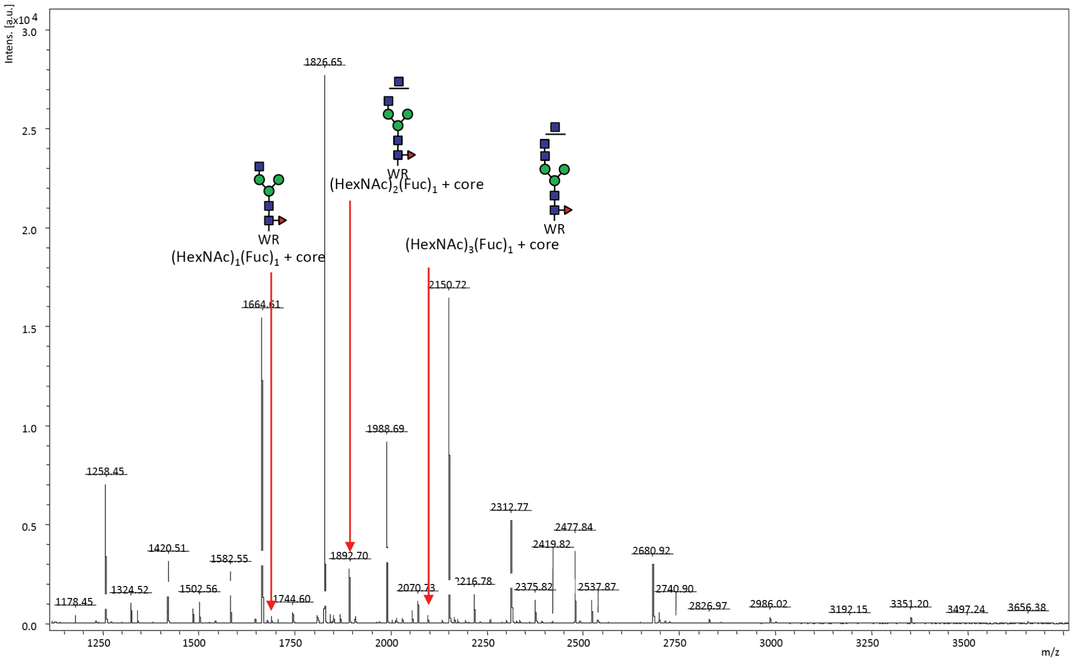


**S4 Figure. Glycoform present in the N-glycans of chondrocytes: core fucose**

N-glycan profiling of chondrocytes isolated from mouse knee cartilage. The released N-glycans using PNGase F are captured and labeled with aoWR using BlotGlyco beads (Sumitomo Bakelite). MALDI-TOF MS analyses of aoWR-labeled glycans were performed using an Autoflex Speed (Bruker Daltonics) operated in positive-ion reflector mode. MALDI-TOF MS, matrix-assisted laser desorption/ionization-time of flight mass spectrometry; aoWR, aminooxy-functionalized tryptophanyl arginine methyl ester.

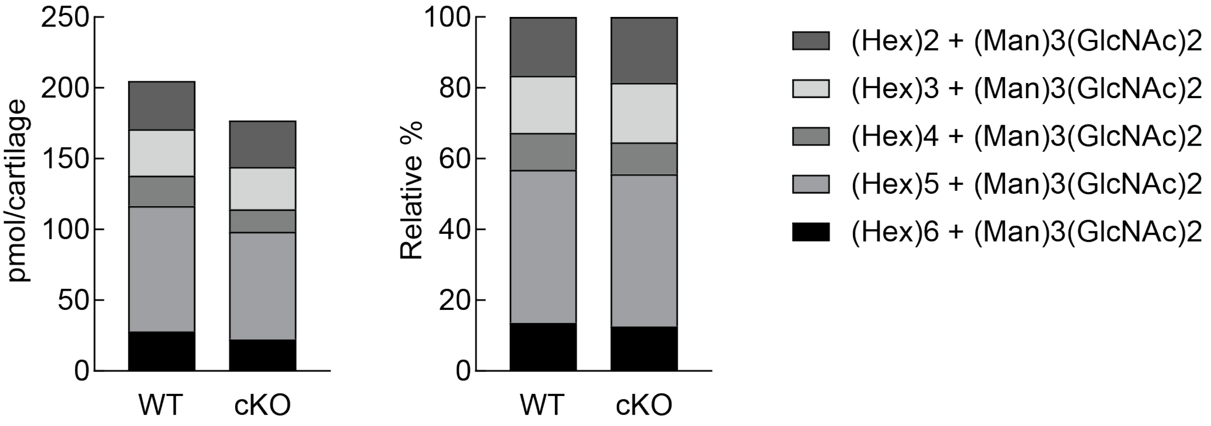


**S5 Figure. Proportion of high-mannose type N-glycans in the cartilage of *Fut8* cKO mice**

Quantified high-mannose type N-glycans (pmol/cartilage) are stratified into HM5 [(Hex)2 + (Man)3(GlcNAc)2], HM6 [(Hex)3 + (Man)3(GlcNAc)2], HM7 [(Hex)4 + (Man)3(GlcNAc)2], HM8 [(Hex)5 + (Man)3(GlcNAc)2], and HM9 [(Hex)6 + (Man)3(GlcNAc)2], according to the glycan structure (left). Comparison of the relative abundance of high-mannose type N-glycans in WT and *Fut8* cKO mouse cartilages (right). WT, wild-type; cKO, conditional knockout.

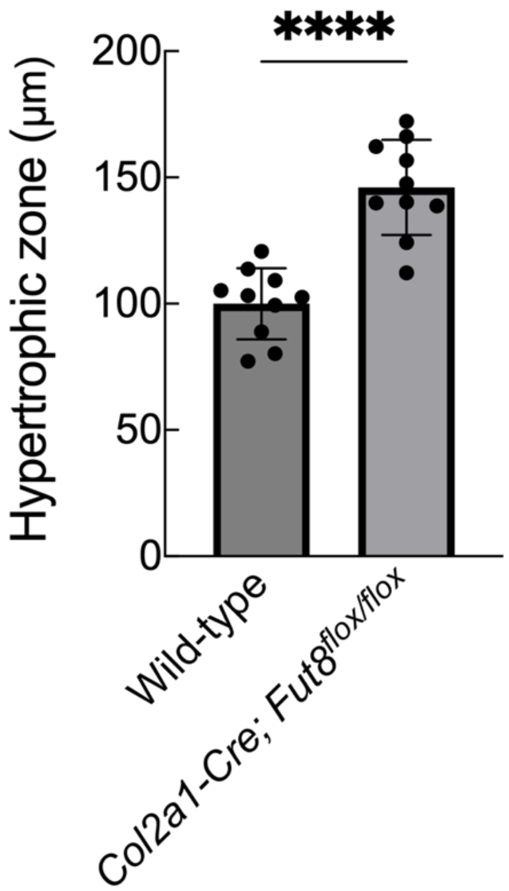


**S6 Figure. Expanded zone of hypertrophic chondrocytes in *Fut8* cKO mice**

Quantification of hypertrophic zones of tibia from 4-week-old WT (C57BL6, 100.1 ± 23.4 µm, n = 10) and *Fut8*$^{-/-}$ (146.1 ± 17.9 µm, n = 10) mice. The length of the hypertrophic zone at the epiphyseal line was measured at three locations (medial, central, and lateral) from stained images of type 10 collagen, and the average of these measurements was representative of the individual. Data are shown as mean ± standard deviation. ****P < 0.0001. The unpaired t-test was used for statistical analysis.

#### S1 Table. N-glycans affected by α-mannosidase

Nineteen of 78 glycans underwent significant changes on either day 3 or 6.

| | Average | | | | SD | | | | *p* -value | |
|---|---|---|---|---|---|---|---|---|---|---|
| | day 3 | | day 6 | | day 3 | | day 6 | | | |
| | man- | man+ | man- | man+ | man- | man+ | man- | man+ | day 3 | day 6 |
| (Hex)2 (HexNAc)2 | 0.792357 | 0.3387174 | 1.6987033 | 0.4567019 | 0.13679 | 0.0508143 | 0.5357954 | 0.0619102 | 0.0057512 | 0.016286 |
| (Hex)2 (HexNAc)2 (Fuc)1 | 3.2347071 | 0.9617929 | 8.2204509 | 0.9248477 | 0.6749618 | 0.1714181 | 2.1522474 | 0.0658934 | 0.0048241 | 0.0042104 |
| (Hex)3 (HexNAc)2 (Fuc)1 | 1.4905489 | 0.8036723 | 2.7099811 | 1.1501296 | 0.2964074 | 0.1192759 | 0.7370968 | 0.1836309 | 0.0204108 | 0.0236565 |
| (Hex)4 (HexNAc)2 | 0.8351768 | 1.5926203 | 1.3871368 | 1.9276786 | 0.1493406 | 0.2261279 | 0.3974899 | 0.2611566 | 0.0083926 | 0.1203717 |
| (Hex)4 (HexNAc)2 (Fuc)1 | 0.3769839 | 0.283877 | 0.5425598 | 0.3520813 | 0.0449464 | 0.0183714 | 0.1063026 | 0.060982 | 0.0293447 | 0.054541 |
| (Hex)2 + (Man)3(GlcNAc)2 | 13.904173 | 17.959473 | 15.158164 | 23.331168 | 1.2342233 | 1.0062143 | 1.7241064 | 1.2449949 | 0.0115924 | 0.0026453 |
| (HexNAc)1 (Fuc)1 + (Man)3(GlcNAc)2 | 1.3015968 | 1.5709037 | 0.815818 | 2.076143 | 0.2070806 | 0.2021715 | 0.0506566 | 0.2089933 | 0.1823063 | 0.0005303 |
| (Hex)2 (Fuc)1 + (Man)3(GlcNAc)2 | 0.322479 | 0.2136506 | 0.4231222 | 0.2682425 | 0.0468934 | 0.0100736 | 0.1128915 | 0.0499029 | 0.0171027 | 0.0954477 |
| (Hex)3 + (Man)3(GlcNAc)2 | 4.8611867 | 5.775936 | 6.6547612 | 5.0794737 | 0.3056028 | 0.3546999 | 1.4178189 | 1.1795482 | 0.0276815 | 0.2131284 |
| (Hex)1 (HexNAc)1 (Fuc)1 + (Man)3(GlcNAc)2 | 0.7502656 | 0.7223147 | 0.5148813 | 0.6088775 | 0.1020876 | 0.016991 | 0.0170882 | 0.0361136 | 0.6642846 | 0.0151606 |
| (HexNAc)2 (Fuc)1 + (Man)3(GlcNAc)2 | 1.652909 | 3.4569757 | 1.3538474 | 3.7150043 | 0.132951 | 0.6544838 | 0.0100189 | 0.3529257 | 0.0094562 | 0.0003174 |
| (Hex)1 (HexNAc)2 + (Man)3(GlcNAc)2 | 0.8761234 | 0.7729216 | 0.6469693 | 0.8172595 | 0.1294936 | 0.0848433 | 0.0886174 | 0.0559445 | 0.3125273 | 0.0480988 |
| (Hex)4 + (Man)3(GlcNAc)2 | 2.4508725 | 2.8951168 | 3.0970917 | 2.2455557 | 0.2531365 | 0.1002396 | 0.766259 | 0.4094851 | 0.0475298 | 0.1648153 |
| (Hex)2 (HexNAc)1 (Fuc)1 + (Man)3(GlcNAc)2 | 0.8195818 | 0.8679034 | 0.6426199 | 0.7280347 | 0.0731323 | 0.0693343 | 0.0472502 | 0.006566 | 0.4529411 | 0.0361773 |
| (HexNAc)3 (Fuc)1 + (Man)3(GlcNAc)2 | 2.0468267 | 2.7447735 | 1.3857347 | 2.4861787 | 0.2568542 | 0.248038 | 0.1225476 | 0.2051412 | 0.0276426 | 0.0013388 |
| (Hex)5 + (Man)3(GlcNAc)2 | 3.3848269 | 2.5793075 | 4.2074717 | 2.518858 | 0.330947 | 0.2363527 | 1.0871818 | 0.3924325 | 0.0265184 | 0.0646356 |
| (HexNAc)4 + (Man)3(GlcNAc)2 | 0.4987153 | 0.3401677 | 0.2835968 | 0.2672303 | 0.0628332 | 0.0278435 | 0.2457312 | 0.2331421 | 0.0161872 | 0.9373258 |
| (Hex)6 + (Man)3(GlcNAc)2 | 6.5672658 | 2.5742614 | 7.8301551 | 3.5422839 | 0.705065 | 0.2276018 | 0.9277843 | 0.2895775 | 0.0007332 | 0.0015756 |
| (Hex)2 (HexNAc)1 (Fuc)1 (NeuAc)1 + (Man)3(GlcNAc)2 | 0.6350732 | 0.5371868 | 0.5159921 | 0.5073008 | 0.0371676 | 0.0445058 | 0.0619954 | 0.0519563 | 0.0430728 | 0.8614184 |

SD; standard deviation (*m/z*/protein)

man-; control, man+; cartilage stimulated by α-mannosidase

#### S2 Table. List of the top 10 deviations from the y = x line

The values are expressed as relative values of the amount of substance (pmol) calculated based on internal standards to the total amount of N-glycan.

| Glycan structure | Control | Mannosidase | Dissociation |
|---|---|---|---|
| (Hex)2 + (Man)3(GlcNAc)2 | 0.13433877 | 0.17518289 | 0.04084413 |
| (HexNAc)2 (Fuc)1 + (Man)3(GlcNAc)2 | 0.01345174 | 0.03953942 | 0.02608767 |
| (Hex)5 + (Man)3(GlcNAc)2 | 0.02873745 | 0.01473236 | 0.01400509 |
| (Hex)6 + (Man)3(GlcNAc)2 | 0.03127904 | 0.01857732 | 0.01270172 |
| (HexNAc)3 (Fuc)1 + (Man)3(GlcNAc)2 | 0.01922672 | 0.03108068 | 0.01185396 |
| (Hex)2 (HexNAc)2 (Fuc)1 | 0.01876579 | 0.00792915 | 0.01083663 |
| (Hex)3 + (Man)3(GlcNAc)2 | 0.03565567 | 0.02670789 | 0.00894778 |
| (HexNAc)1 (Fuc)1 + (Man)3(GlcNAc)2 | 0.01320084 | 0.01968082 | 0.00647997 |
| (HexNAc)4 (Fuc)2 + (Man)3(GlcNAc)2 | 0.05688225 | 0.05042987 | 0.00645238 |

p mol relative (%)

### S3 Table. List of glycans quantified in healthy and osteoarthritic human cartilage

**N-glycan**

| No. | Glycan compositioin | Class | Theor *m/z* | sample (pmol/100 µg protein) | | | | | | | | | | Average | |
|---|---|---|---|---|---|---|---|---|---|---|---|---|---|---|---|
| | | | | Ctrl-1 | Ctrl-2 | Ctrl-3 | Ctrl-4 | Ctrl-5 | OA-1 | OA-2 | OA-3 | OA-4 | OA-5 | Ctrl | OA |
| 1 | (Hex)3 (HexNAc)2 | | 1340.55 | 4.5 | 0.0 | 6.3 | 0.0 | 4.3 | 0.0 | 0.0 | 0.0 | 0.0 | 0.0 | 3.0 | 0.0 |
| 2 | (Hex)4 (HexNAc)2 | | 1502.60 | 8.9 | 7.2 | 16.6 | 9.4 | 6.5 | 0.0 | 10.2 | 3.6 | 0.0 | 0.0 | 9.7 | 2.8 |
| 3 | (Hex)2 + (Man)3(GlcNAc)2 | | 1664.66 | 750.2 | 499.2 | 1121.8 | 889.7 | 558.9 | 147.7 | 792.0 | 242.0 | 508.6 | 264.2 | 764.0 | 390.9 |
| 4 | (Hex)3 + (Man)3(GlcNAc)2 | High mannose (HM) | 1826.71 | 640.3 | 338.6 | 405.2 | 631.2 | 426.8 | 95.8 | 396.9 | 320.9 | 259.5 | 267.5 | 488.4 | 268.1 |
| 5 | (Hex)4 + (Man)3(GlcNAc)2 | | 1988.76 | 28.6 | 10.6 | 15.9 | 20.8 | 11.4 | 9.2 | 30.2 | 20.7 | 21.8 | 15.9 | 17.4 | 19.6 |
| 6 | (Hex)5 + (Man)3(GlcNAc)2 | | 2150.81 | 12.0 | 6.7 | 11.4 | 9.4 | 0.0 | 5.7 | 32.6 | 13.0 | 19.7 | 11.2 | 7.9 | 16.4 |
| 7 | (Hex)6 + (Man)3(GlcNAc)2 | | 2312.87 | 9.5 | 9.2 | 13.5 | 14.0 | 0.0 | 0.0 | 24.7 | 12.4 | 26.1 | 13.8 | 9.2 | 15.4 |
| 8 | (HexNAc)1 + (Man)3(GlcNAc)2 | | 1543.63 | 6.9 | 6.5 | 11.9 | 6.2 | 6.8 | 5.3 | 11.5 | 7.0 | 38.3 | 9.2 | 7.7 | 14.3 |
| 9 | (HexNAc)1 (Fuc)1 + (Man)3(GlcNAc)2 | | 1689.69 | 19.9 | 12.7 | 23.6 | 15.7 | 17.1 | 9.0 | 29.5 | 15.1 | 39.2 | 14.0 | 17.8 | 21.4 |
| 10 | (Hex)1 (HexNAc)1 + (Man)3(GlcNAc)2 | | 1705.68 | 7.7 | 4.9 | 7.6 | 5.6 | 4.0 | 0.0 | 13.1 | 5.0 | 29.9 | 8.7 | 5.9 | 11.3 |
| 11 | (HexNAc)2(Fuc)1+(Man)2(GlcNAc)2 | | 1730.71 | 0.0 | 0.0 | 0.0 | 3.9 | 0.0 | 0.0 | 0.0 | 0.0 | 0.0 | 0.0 | 0.8 | 0.0 |
| 12 | (HexNAc)2 + (Man)3(GlcNAc)2 | | 1746.71 | 7.8 | 8.4 | 8.2 | 6.0 | 3.9 | 2.5 | 14.4 | 4.2 | 19.5 | 8.8 | 6.9 | 9.9 |
| 13 | (Hex)1 (HexNAc)1 (Fuc)1 + (Man)3(GlcNAc)2 | | 1851.74 | 14.8 | 9.6 | 16.5 | 13.6 | 12.3 | 8.0 | 25.6 | 11.0 | 35.5 | 14.4 | 13.4 | 18.9 |
| 14 | (Hex)2 (HexNAc)1 + (Man)3(GlcNAc)2 | | 1867.74 | 7.8 | 4.7 | 5.6 | 5.0 | 5.0 | 0.0 | 10.2 | 5.7 | 15.8 | 7.1 | 5.6 | 7.8 |
| 15 | (HexNAc)2 (Fuc)1 + (Man)3(GlcNAc)2 | | 1892.77 | 32.3 | 43.7 | 39.8 | 21.3 | 19.4 | 14.3 | 44.7 | 25.0 | 78.2 | 36.8 | 31.3 | 39.8 |
| 16 | (Hex)1 (HexNAc)2 + (Man)3(GlcNAc)2 | | 1908.76 | 11.3 | 7.9 | 12.7 | 9.1 | 7.4 | 0.0 | 26.1 | 10.4 | 51.4 | 11.8 | 9.7 | 19.9 |
| 17 | (HexNAc)3 + (Man)3(GlcNAc)2 | | 1949.79 | 0.0 | 3.9 | 4.1 | 0.0 | 0.0 | 0.0 | 6.9 | 0.0 | 0.0 | 0.0 | 1.6 | 1.4 |
| 18 | (Hex)3 (HexNAc)1 + (Man)3(GlcNAc)2 | | 2029.79 | 8.4 | 0.0 | 0.0 | 6.7 | 4.6 | 0.0 | 10.3 | 4.7 | 0.0 | 4.7 | 3.9 | 3.9 |
| 19 | (HexNAc)2(Fuc)2+ (Man)3(GlcNAc)2 | | 2038.82 | 50.9 | 40.5 | 39.0 | 19.2 | 19.7 | 17.1 | 40.3 | 24.3 | 42.1 | 47.3 | 33.9 | 34.2 |
| 20 | (Hex)1 (HexNAc)2 (Fuc)1 + (Man)3(GlcNAc)2 | | 2054.82 | 72.3 | 52.0 | 69.7 | 42.7 | 46.9 | 31.8 | 133.0 | 62.6 | 248.0 | 65.9 | 56.7 | 108.3 |
| 21 | (Hex)2 (HexNAc)2 + (Man)3(GlcNAc)2 | Neutral complex/ hybrid (C/H-N) | 2070.81 | 8.9 | 6.0 | 8.2 | 8.4 | 5.4 | 8.0 | 17.1 | 9.7 | 51.4 | 8.4 | 7.4 | 18.9 |
| 22 | (HexNAc)3 (Fuc)1 + (Man)3(GlcNAc)2 | | 2095.85 | 23.4 | 21.4 | 24.8 | 20.8 | 13.5 | 9.7 | 82.9 | 24.3 | 51.1 | 20.4 | 20.8 | 37.7 |
| 23 | (Hex)1 (HexNAc)3 + (Man)3(GlcNAc)2 | | 2111.84 | 6.2 | 5.7 | 8.9 | 6.3 | 4.9 | 5.9 | 14.3 | 9.6 | 18.0 | 6.7 | 6.4 | 10.9 |
| 24 | (HexNAc)4 + (Man)3(GlcNAc)2 | | 2152.87 | 0.0 | 0.0 | 0.0 | 3.9 | 0.0 | 0.0 | 7.6 | 4.6 | 0.0 | 0.0 | 0.8 | 2.4 |
| 25 | (Hex)1 (HexNAc)2 (Fuc)2 + (Man)3(GlcNAc)2 | | 2200.88 | 31.6 | 15.4 | 29.3 | 23.4 | 24.9 | 26.4 | 47.8 | 33.7 | 39.8 | 35.9 | 24.9 | 36.7 |
| 26 | (Hex)2 (HexNAc)2 (Fuc)1+ (Man)3(GlcNAc)2 | | 2216.87 | 99.3 | 43.1 | 61.6 | 58.8 | 49.7 | 49.9 | 109.3 | 95.4 | 340.9 | 58.7 | 62.5 | 130.9 |
| 27 | (HexNAc)3 (Fuc)2+ (Man)3(GlcNAc)2 | | 2241.90 | 25.0 | 20.7 | 30.8 | 15.7 | 19.4 | 0.0 | 19.9 | 12.8 | 20.8 | 10.2 | 22.3 | 12.7 |
| 28 | (Hex)1 (HexNAc)3 (Fuc)1 + (Man)3(GlcNAc)2 | | 2257.90 | 64.7 | 42.9 | 66.4 | 61.9 | 42.9 | 36.6 | 336.7 | 105.9 | 186.3 | 71.7 | 55.8 | 147.5 |
| 29 | (HexNAc)4 (Fuc)1 + (Man)3(GlcNAc)2 | | 2298.93 | 22.3 | 17.1 | 29.3 | 20.2 | 18.0 | 13.3 | 52.9 | 23.3 | 55.9 | 20.7 | 21.4 | 33.2 |
| 30 | (Hex)2 (HexNAc)2(Fuc)2 + (Man)3(GlcNAc)2 | | 2362.93 | 7.2 | 4.6 | 7.0 | 6.2 | 5.6 | 9.0 | 12.5 | 10.2 | 12.0 | 6.9 | 6.1 | 10.1 |
| 31 | (Hex)1 (HexNAc)3(Fuc)2 + (Man)3(GlcNAc)2 | | 2403.96 | 137.4 | 67.1 | 94.4 | 66.3 | 66.7 | 53.1 | 185.3 | 113.4 | 126.0 | 63.4 | 86.4 | 108.2 |
| 32 | (Hex)2 (HexNAc)3 (Fuc)1 + (Man)3(GlcNAc)2 | | 2419.95 | 7.6 | 5.1 | 5.1 | 8.5 | 5.0 | 5.7 | 31.6 | 10.6 | 11.7 | 7.2 | 6.3 | 13.4 |
| 33 | (HexNAc)4 (Fuc)2 + (Man)3(GlcNAc)2 | | 2444.98 | 198.6 | 118.1 | 208.2 | 131.7 | 131.9 | 68.3 | 213.4 | 142.5 | 127.5 | 99.6 | 157.7 | 130.3 |
| 34 | (Hex)3 (HexNAc)3 (Fuc)1 + (Man)3(GlcNAc)2 | | 2582.00 | 4.8 | 3.9 | 4.8 | 3.9 | 4.2 | 0.0 | 6.2 | 5.5 | 0.0 | 0.0 | 4.3 | 2.3 |
| 35 | (HexNAc)4 (Fuc)3 + (Man)3(GlcNAc)2 | | 2591.04 | 216.2 | 44.5 | 75.7 | 124.7 | 131.9 | 69.2 | 172.1 | 153.1 | 97.5 | 86.2 | 118.6 | 115.6 |
| 36 | (HexNAc)2 (NeuAc)1[a2,6]+ (Man)3(GlcNAc)2 | | 2078.87 | 0.0 | 0.0 | 4.4 | 0.0 | 0.0 | 0.0 | 0.0 | 0.0 | 0.0 | 0.0 | 0.9 | 0.0 |
| 37 | (Hex)1 (HexNAc)1 (Fuc)1 (NeuAc)1[a2,6]+ (Man)3(GlcNAc)2 | | 2183.90 | 0.0 | 0.0 | 0.0 | 0.0 | 3.1 | 0.0 | 0.0 | 0.0 | 0.0 | 0.0 | 0.6 | 0.0 |
| 38 | (HexNAc)2 (Fuc)1 (NeuAc)1[a2,6] + (Man)3(GlcNAc)2 | | 2224.93 | 5.9 | 5.1 | 8.7 | 0.0 | 3.1 | 0.0 | 6.2 | 0.0 | 0.0 | 5.7 | 4.5 | 2.4 |
| 39 | (Hex)1 (HexNAc)2 (NeuAc)1[a2,6] + (Man)3(GlcNAc)2 | | 2240.92 | 0.0 | 0.0 | 0.0 | 0.0 | 0.0 | 7.0 | 0.0 | 0.0 | 0.0 | 0.0 | 0.0 | 1.4 |
| 40 | (HexNAc)3(NeuAc)1[a2,6] + (Man)3(GlcNAc)2 | | 2281.95 | 3.9 | 0.0 | 0.0 | 0.0 | 0.0 | 0.0 | 0.0 | 0.0 | 0.0 | 0.0 | 0.8 | 0.0 |
| 41 | (Hex)1 (HexNAc)2 (Fuc)1 (NeuAc)1[a2,3] + (Man)3(GlcNAc)2 | | 2358.94 | 4.3 | 5.0 | 0.0 | 0.0 | 0.0 | 0.0 | 0.0 | 0.0 | 0.0 | 0.0 | 1.9 | 0.0 |
| 42 | (Hex)1 (HexNAc)2 (Fuc)1 (NeuAc)1[a2,6] + (Man)3(GlcNAc)2 | | 2386.98 | 33.4 | 28.0 | 27.4 | 15.9 | 19.5 | 6.9 | 14.6 | 13.7 | 18.5 | 10.0 | 24.8 | 12.7 |
| 43 | (HexNAc)3 (Fuc)1 (NeuAc)1[a2,6] + (Man)3(GlcNAc)2 | | 2428.00 | 17.7 | 16.4 | 30.6 | 10.5 | 11.1 | 0.0 | 9.9 | 6.1 | 0.0 | 5.9 | 17.3 | 4.4 |
| 44 | (HexNAc)4 (NeuAc)1[a2,6] + (Man)3(GlcNAc)2 | | 2485.03 | 4.8 | 0.0 | 6.8 | 3.9 | 2.8 | 0.0 | 0.0 | 0.0 | 0.0 | 0.0 | 3.7 | 0.0 |
| 45 | (Hex)2 (HexNAc)2 (Fuc)1 (NeuAc)1[a2,3] + (Man)3(GlcNAc)2 | Sialylated complex/ hybrid (C/H-N) | 2521.00 | 23.5 | 11.5 | 9.0 | 6.4 | 3.5 | 10.3 | 10.4 | 14.4 | 26.9 | 7.4 | 10.8 | 13.9 |
| 46 | (Hex)2 (HexNAc)2 (Fuc)1 (NeuAc)1[a2,6] + (Man)3(GlcNAc)2 | | 2549.03 | 311.7 | 146.0 | 105.4 | 179.7 | 123.5 | 54.2 | 143.1 | 119.8 | 172.8 | 62.0 | 173.3 | 110.4 |
| 47 | (Hex)1 (HexNAc)3 (Fuc)1 (NeuAc)1[a2,6] + (Man)3(GlcNAc)2 | | 2590.06 | 143.1 | 151.9 | 262.9 | 100.1 | 75.4 | 16.0 | 95.8 | 45.4 | 41.9 | 29.5 | 146.7 | 45.7 |
| 48 | (HexNAc)4(Fuc)1(NeuAc)1[a2,6] + (Man)3(GlcNAc)2 | | 2631.08 | 38.2 | 30.1 | 62.4 | 37.6 | 28.3 | 17.3 | 0.0 | 22.6 | 24.0 | 15.1 | 39.3 | 15.8 |
| 49 | (Hex)1 (HexNAc)3 (Fuc)2 (NeuAc)1[a2,3] + (Man)3(GlcNAc)2 | | 2708.08 | 0.0 | 3.7 | 3.8 | 0.0 | 0.0 | 0.0 | 0.0 | 5.4 | 0.0 | 0.0 | 1.5 | 1.1 |
| 50 | (Hex)1(HexNAc)3 (Fuc)2 (NeuAc)1[a2,6] + (Man)3(GlcNAc)2 | | 2736.12 | 10.8 | 9.0 | 9.2 | 5.1 | 4.6 | 0.0 | 7.6 | 9.5 | 9.8 | 0.0 | 7.8 | 5.4 |
| 51 | (HexNAc)4 (Fuc)2 (NeuAc)1[a2,6] + (Man)3(GlcNAc)2 | | 2777.14 | 14.6 | 8.5 | 16.3 | 9.4 | 7.9 | 5.9 | 7.9 | 7.5 | 0.0 | 4.3 | 11.3 | 5.1 |
| 52 | (Hex)2 (HexNAc)2 (Fuc)1 (NeuAc)2[a2,3/a2,6] + (Man)3(GlcNAc)2 | | 2853.16 | 7.8 | 4.1 | 4.9 | 2.6 | 0.0 | 0.0 | 0.0 | 4.2 | 0.0 | 0.0 | 3.9 | 0.8 |
| 53 | (Hex)1 (HexNAc)3 (Fuc)1 (NeuAc)2[a2,3/a2,6] + (Man)3(GlcNAc)2 | | 2894.18 | 0.0 | 3.2 | 5.8 | 0.0 | 0.0 | 0.0 | 0.0 | 0.0 | 0.0 | 0.0 | 1.8 | 0.0 |
| 54 | (Hex)3 (HexNAc)3 (Fuc)1 (NeuAc)1[a2,6] + (Man)3(GlcNAc)2 | | 2914.16 | 7.5 | 4.3 | 0.0 | 0.0 | 2.0 | 0.0 | 0.0 | 0.0 | 0.0 | 0.0 | 2.7 | 0.0 |
| | Total amount of *N*-glycan pmol/µg | | | 3175.0 | 1908.3 | 3041.8 | 2661.2 | 1963.9 | 819.2 | 3257.2 | 1790.6 | 2866.4 | 1437.2 | 2550.0 | 2034.1 |

**O-glycan**

| No. | Glycan composition | Class | Theor m/z | Sample | | | | | | | | | | Average | |
|---|---|---|---|---|---|---|---|---|---|---|---|---|---|---|---|
| | | | | Ctrl-1 | Ctrl-2 | Ctrl-3 | Ctrl-4 | Ctrl-5 | OA-1 | OA-2 | OA-3 | OA-4 | OA-5 | Ctrl | OA |
| 1 | (HexNAc)1 | Core1 | 551.383 | 360.93 | 237.13 | 293.57 | 112.18 | 175.03 | 204.15 | 544.80 | 283.60 | 314.91 | 247.17 | 235.77 | 318.92 |
| 2 | (HexNAc)1(Hex)1 | Core1 | 713.614 | 101.84 | 87.65 | 103.86 | 70.84 | 82.55 | 49.04 | 177.56 | 93.52 | 81.21 | 52.24 | 89.35 | 90.71 |
| 3 | (HexNAc)1(Neu5Ac)1 | Core1 | 842.744 | 10.62 | 9.32 | 4.95 | 2.11 | 3.61 | 5.75 | 10.03 | 5.23 | 7.30 | 3.42 | 6.12 | 6.35 |
| 4 | (HexNAc)1(Hex)1(Neu5Ac)1 | Core1 | 1004.917 | 138.06 | 90.82 | 120.64 | 65.12 | 90.31 | 45.55 | 120.01 | 85.70 | 84.80 | 36.40 | 100.99 | 74.49 |
| 5 | (HexNAc)1(Hex)1(Neu5Ac)1 Na+ | | 1026.887 | | | | | | | | | | | | |
| 6 | (HexNAc)1(Hex)1(Neu5Ac)2 | Core1 | 1296.172 | 47.88 | 53.09 | 25.83 | 16.58 | 24.19 | 24.05 | 32.86 | 28.72 | 28.41 | 6.74 | 33.51 | 24.16 |
| 7 | (HexNAc)1(Hex)1(Neu5Ac)2 Na+ | | 1318.167 | | | | | | | | | | | | |
| 8 | (HexNAc)2(Hex)1 | Core2 | 916.847 | 14.01 | 13.90 | 16.63 | 15.18 | 13.86 | 4.68 | 23.82 | 16.05 | 6.48 | 6.54 | 14.72 | 11.51 |
| 9 | (HexNAc)2(Hex)2 | Core2 | 1079.009 | 81.69 | 51.20 | 63.71 | 88.96 | 64.15 | 25.46 | 137.23 | 72.57 | 28.16 | 36.87 | 69.94 | 60.06 |
| 10 | (HexNAc)2(Hex)2 Na+ | | 1100.944 | | | | | | | | | | | | |
| 11 | (HexNAc)2(Hex)1(Neu5Ac)1 | Core2 | 1208.096 | 4.77 | 4.92 | 5.93 | 3.55 | 2.95 | 2.42 | 3.55 | 3.29 | 1.51 | 1.20 | 4.42 | 2.39 |
| 12 | (HexNAc)2(Hex)2(Neu5Ac)1 | Core2 | 1370.255 | 52.97 | 43.18 | 67.73 | 52.56 | 37.00 | 17.91 | 44.16 | 40.60 | 11.94 | 11.40 | 50.69 | 25.20 |
| 13 | (HexNAc)2(Hex)2(Neu5Ac)1 Na+ | | 1392.218 | | | | | | | | | | | | |
| 14 | (HexNAc)2(Hex)2(Neu5Ac)2 | Core2 | 1661.498 | 12.34 | 11.54 | 14.30 | 6.65 | 5.39 | 9.89 | 4.76 | 6.35 | 3.80 | 1.81 | 10.04 | 5.32 |
| 15 | (HexNAc)2 | Core3 | 754.666 | 39.71 | 16.37 | 18.17 | 27.39 | 17.77 | 15.23 | 57.52 | 26.64 | 11.91 | 18.49 | 23.88 | 25.96 |
| 16 | (Xyl)1 | GAG | 480.288 | 34.82 | 56.97 | 17.44 | 7.38 | 12.05 | 16.07 | 25.09 | 25.99 | 19.76 | 26.06 | 25.73 | 22.59 |
| 17 | (Hex)1(Xyl)1 | GAG | 642.496 | 5.41 | 8.86 | 2.77 | 1.71 | 1.81 | 3.37 | 5.43 | 4.16 | 2.87 | 4.26 | 4.11 | 4.02 |
| 18 | (Hex)2(Xyl)1 | GAG | 804.707 | 3.38 | 2.85 | 0.00 | 0.00 | 0.00 | 0.00 | 0.00 | 0.00 | 0.00 | 1.47 | 1.25 | 0.29 |
| 19 | (Hex)1(Xyl)1(NeuAc)1 | GAG | 933.745 | 5.62 | 36.05 | 0.00 | 0.58 | 0.00 | 0.00 | 0.00 | 2.30 | 0.00 | 3.10 | 8.45 | 1.08 |
| 20 | (GluA)1(Hex)2(Xyl)1 | GAG | 980.863 | 2.03 | 2.68 | 1.32 | 1.23 | 1.33 | 0.00 | 2.11 | 2.04 | 1.40 | 2.01 | 1.72 | 1.51 |
| 21 | (NeuAc)1(Hex)2(Xyl)1 | GAG | 1095.988 | 1.98 | 0.91 | 1.07 | 0.00 | 0.00 | 0.00 | 0.00 | 0.00 | 0.00 | 0.00 | 0.79 | 0.00 |
| 22 | (HexNAc)1(GlcA)1(Hex)2(Xly)1 | GAG | 1184.054 | 13.03 | 9.78 | 6.33 | 9.01 | 10.81 | 3.61 | 5.67 | 8.55 | 4.52 | 4.69 | 9.79 | 5.41 |
| | Total amount of *O*-glycan | | | 931.1 | 737.2 | 764.3 | 481.0 | 542.8 | 427.2 | 1194.6 | 705.3 | 609.0 | 463.9 | 691.3 | 680.0 |

**GSL-glycan**

| No. | Glycan composition | Class | Theor m/z | Sample | | | | | | | | | | Average | |
|---|---|---|---|---|---|---|---|---|---|---|---|---|---|---|---|
| | | | | Ctrl-1 | Ctrl-2 | Ctrl-3 | Ctrl-4 | Ctrl-5 | OA-1 | OA-2 | OA-3 | OA-4 | OA-5 | Ctrl | OA |
| 1 | (Hex)2 | Lac | 772.336 | 182.72 | 701.76 | 1456.10 | 195.81 | 350.22 | 483.48 | 180.71 | 387.65 | 216.29 | 1551.36 | 577.32 | 563.90 |
| 2 | (Hex)3 | Gb | 934.389 | 80.71 | 392.96 | 671.56 | 21.86 | 134.18 | 386.51 | 81.05 | 253.06 | 0.00 | 1153.41 | 260.25 | 374.81 |
| 9 | (Hex)4 | Gb | 1096.442 | 0.00 | 80.44 | 361.50 | 0.00 | 22.54 | 149.60 | 7.04 | 75.82 | 0.00 | 1238.57 | 92.90 | 294.20 |
| 15 | (Hex)5 | Gb | 1258.495 | 0.00 | 0.00 | 0.00 | 0.00 | 0.00 | 0.00 | 0.00 | 0.00 | 0.00 | 120.43 | 0.00 | 24.09 |
| 27 | (Hex)6 | Gb | 1420.548 | 0.00 | 0.00 | 0.00 | 0.00 | 0.00 | 0.00 | 0.00 | 0.00 | 0.00 | 44.21 | 0.00 | 8.84 |
| 41 | (Hex)7 | Gb | 1582.600 | 0.00 | 0.00 | 81.94 | 0.00 | 0.00 | 0.00 | 0.00 | 0.00 | 0.00 | 28.45 | 16.39 | 5.69 |
| 3 | (Hex)2(HexNAc)1 | (n)Lc | 975.416 | 0.00 | 0.00 | 0.00 | 90.97 | 0.00 | 0.00 | 0.00 | 0.00 | 0.00 | 0.00 | 18.19 | 0.00 |
| 10 | (Hex)3(HexNAc)1 | (n)Lc | 1137.469 | 886.12 | 2361.98 | 3421.95 | 975.57 | 1646.27 | 3703.84 | 2222.90 | 3604.64 | 1285.06 | 6157.77 | 1858.38 | 3394.84 |
| 23 | (Hex)3(HexNAc)2 | (n)Lc | 1340.548 | 0.00 | 0.00 | 0.00 | 0.00 | 43.05 | 0.00 | 0.00 | 0.00 | 0.00 | 0.00 | 8.61 | 0.00 |
| 36 | (Hex)4(HexNAc)2 | (n)Lc | 1502.601 | 9.64 | 0.00 | 144.32 | 70.32 | 73.13 | 0.00 | 0.00 | 20.20 | 0.00 | 0.00 | 59.48 | 4.04 |
| 4 | (Hex)2(a2,3NeuAc)1 | Gg | 1076.464 | 255.91 | 1556.47 | 1777.79 | 336.33 | 408.84 | 861.76 | 175.55 | 440.02 | 242.61 | 1779.40 | 867.07 | 699.87 |
| 24 | (Hex)2(a2,3/a2,3NeuAc)2 | Gg | 1380.591 | 197.94 | 598.40 | 204.82 | 166.53 | 134.44 | 290.02 | 114.77 | 171.94 | 85.54 | 660.93 | 260.43 | 264.64 |
| 29 | (Hex)3(HexNAc)1(a2,3NeuAc)1 | Gg | 1441.596 | 139.47 | 419.95 | 1638.09 | 296.21 | 494.01 | 733.72 | 390.79 | 568.19 | 105.49 | 1095.07 | 597.55 | 578.65 |
| 30 | (Hex)3(HexNAc)1(a2,6NeuAc)1 | Gg | 1469.626 | 19.84 | 51.55 | 91.58 | 69.69 | 56.25 | 62.16 | 24.18 | 34.54 | 0.00 | 102.66 | 57.78 | 44.71 |
| 59 | (Hex)2(a2,3NeuAc)3 | Gg | 1684.718 | 10.51 | 17.49 | 0.00 | 13.46 | 0.00 | 0.00 | 0.00 | 0.00 | 0.00 | 0.00 | 8.29 | 0.00 |
| 72 | (Hex)3(HexNAc)1(a2,3NeuAc)2 | Gg | 1745.723 | 27.19 | 91.54 | 736.16 | 81.96 | 131.11 | 206.80 | 140.18 | 146.13 | 16.92 | 337.40 | 213.59 | 169.49 |
| 88 | (Hex)4(HexNAc)2(a2,3NeuAc)1 | Gg | 1806.728 | 15.50 | 28.82 | 71.73 | 64.80 | 72.78 | 34.39 | 28.65 | 44.94 | 0.00 | 57.76 | 50.73 | 33.15 |
| 139 | (Hex)3(HexNAc)1(a2,3NeuAc)3 | Gg | 2049.850 | 3.09 | 10.93 | 78.53 | 10.51 | 10.54 | 25.44 | 32.14 | 14.56 | 0.00 | 34.68 | 22.72 | 21.37 |
| | Total amount of *GSL*-glycan | | | 1828.6 | 6312.3 | 10736.1 | 2394.0 | 3577.3 | 6937.7 | 3398.0 | 5761.7 | 1951.9 | 14362.1 | 4969.7 | 6482.3 |

**GAG**

| No. | Glycan composition | Class | Sample | | | | | | | | | | Average | |
|---|---|---|---|---|---|---|---|---|---|---|---|---|---|---|
| | | | Ctrl-1 | Ctrl-2 | Ctrl-3 | Ctrl-4 | Ctrl-5 | OA-1 | OA-2 | OA-3 | OA-4 | OA-5 | Ctrl | OA |
| 1 | ΔHA | HA | 24.71 | 34.01 | 22.43 | 44.45 | 36.27 | 8.02 | 17.99 | 13.17 | 12.56 | 11.11 | 32.37 | 12.57 |
| 2 | ΔCS-CS0S | CS | 11.54 | 8.61 | 6.12 | 7.60 | 7.39 | 14.55 | 20.94 | 31.87 | 34.76 | 20.23 | 8.25 | 24.47 |
| 3 | ΔCS-CS4S | CS | 36.10 | 55.06 | 29.43 | 43.53 | 38.98 | 35.01 | 54.29 | 42.31 | 37.98 | 48.24 | 40.62 | 43.57 |
| 4 | ΔCS-CS6S | CS | 187.56 | 209.78 | 180.25 | 171.33 | 135.87 | 78.91 | 503.89 | 354.61 | 177.98 | 193.90 | 176.96 | 261.86 |
| 5 | ΔCS-CS2S6S | CS | 1.83 | 4.88 | 2.76 | 2.66 | 1.44 | 1.82 | 7.87 | 4.71 | 1.79 | 5.54 | 2.71 | 4.34 |
| 6 | ΔCS-CS2S4S | CS | 0.15 | 1.03 | 0.10 | 0.00 | 0.00 | 0.00 | 1.32 | 1.15 | 0.25 | 0.00 | 0.26 | 0.54 |
| 7 | ΔCS-CS4S6S | CS | 0.32 | 0.00 | 0.00 | 0.33 | 0.20 | 0.35 | 0.61 | 1.38 | 0.40 | 0.00 | 0.17 | 0.55 |
| 8 | ΔHS-0S | HS | 3.93 | 3.15 | 3.57 | 3.24 | 3.06 | 2.79 | 2.27 | 3.03 | 4.68 | 4.86 | 3.39 | 3.53 |
| 9 | ΔHS-6S | HS | 0.34 | 0.51 | 0.41 | 0.00 | 0.40 | 0.38 | 0.29 | 0.37 | 0.28 | 0.00 | 0.33 | 0.26 |
| 10 | ΔHS-NS | HS | 1.16 | 2.50 | 1.61 | 1.36 | 1.61 | 2.50 | 3.12 | 4.33 | 1.03 | 2.46 | 1.65 | 2.69 |
| 11 | ΔHS-2S6S | HS | 0.00 | 0.13 | 0.00 | 0.43 | 0.00 | 0.00 | 0.00 | 0.00 | 0.00 | 0.00 | 0.11 | 0.00 |
| 12 | ΔHS-2SNS | HS | 0.00 | 0.00 | 0.00 | 0.19 | 0.00 | 0.17 | 0.43 | 1.44 | 0.10 | 0.00 | 0.04 | 0.43 |
| 13 | ΔHS-NS6S | HS | 0.00 | 0.00 | 0.00 | 0.00 | 0.00 | 0.07 | 0.18 | 0.65 | 0.00 | 0.00 | 0.00 | 0.18 |
| | Total amount of *GAG* | | 267.628 | 319.7 | 246.7 | 275.1 | 225.2 | 144.6 | 613.2 | 459.0 | 271.8 | 286.4 | 266.9 | 355.0 |

**fOS**

| No. | Glycan composition | Class | Theor m/z | Sample | | | | | | | | | | Average | |
|---|---|---|---|---|---|---|---|---|---|---|---|---|---|---|---|
| | | | | Ctrl-1 | Ctrl-2 | Ctrl-3 | Ctrl-4 | Ctrl-5 | OA-1 | OA-2 | OA-3 | OA-4 | OA-5 | Ctrl | OA |
| 1 | Hex1 | | 610.28 | 31.6 | 0.0 | 73.9 | 457.1 | 44.0 | 21.0 | 94.1 | 43.5 | 131.9 | 91.8 | 121.3 | 76.4 |
| 2 | Hex3 | | 934.39 | 0.0 | 0.0 | 98.9 | 56.6 | 0.0 | 19.9 | 40.4 | 144.5 | 761.5 | 355.2 | 31.1 | 264.3 |
| 3 | Hex4 | Hexose oligoner | 1096.44 | 16.4 | 11.2 | 134.8 | 76.1 | 23.9 | 42.2 | 40.9 | 247.7 | 1302.6 | 620.6 | 52.5 | 450.8 |
| 4 | Hex5 | | 1258.49 | 0.0 | 0.0 | 0.0 | 0.0 | 0.0 | 5.6 | 0.0 | 0.0 | 59.7 | 46.2 | 0.0 | 22.3 |
| 5 | Hex6 | | 1420.55 | 0.0 | 0.0 | 0.0 | 0.0 | 5.7 | 4.7 | 0.0 | 0.0 | 25.4 | 19.0 | 1.1 | 9.8 |
| 6 | Hex7 | | 1582.60 | 0.0 | 0.0 | 0.0 | 0.0 | 4.6 | 0.0 | 0.0 | 0.0 | 0.0 | 16.6 | 0.9 | 3.3 |
| 7 | (HexNAc)1 | N-glycan related fOS | 651.31 | 242.2 | 55.2 | 587.3 | 141.6 | 212.7 | 97.3 | 169.8 | 177.8 | 558.8 | 643.9 | 247.8 | 329.5 |
| 8 | (Hex)2 (HexNAc)1 | | 975.42 | 12.8 | 7.7 | 0.0 | 0.0 | 0.0 | 0.0 | 0.0 | 34.0 | 0.0 | 0.0 | 4.1 | 6.8 |
| | Total amount of *fOS* | | | 303.0 | 74.1 | 894.9 | 731.4 | 291.0 | 190.6 | 345.1 | 647.4 | 2840.0 | 1793.4 | 458.9 | 1163.3 |

**S4 Table. Primers used in the gene expression experiment**

The values are expressed as relative values of the amount of substance (pmol) calculated based on internal standards to the total amount of N-glycan.

| Gene | | Sequence |
|---|---|---|
| *Ppia* | Forward | GGTGGTGACTTCACACGCCATAATG |
| | Reverse | CTTGCCATCCAACCACTCAGTCTTG |
| *Fut8* | Forward | TGTTCTGGCTGAGGCCATCTA |
| | Reverse | GTGCCCAGAGTAACAGCAGGA |
| *Tgf-β* | Forward | CTGCTGACCCCCACTGATAC |
| | Reverse | AGCCCTGTATTCCGTCTCCT |
| *Mmp13* | Forward | TTGGCCACTCCCTAGGTCTG |
| | Reverse | GGTTGGGGTCTTCATCGC |
| *Col10a1* | Forward | GGCTTCAGGGAGTGCAATC |
| | Reverse | CTCACATGGGAGCCACTAGG |
| *Col2a1* | Forward | AGGATGGCTGCACCAAACAC |
| | Reverse | TGTCCATGGGTGCGATGTC |
| *Ihh* | Forward | CTCTTGCCTACAAGCAGTTCA |
| | Reverse | CCGTGTTCTCCTCGTCCTT |
| *Aggrecan* | Forward | CCCTCACCCCAAGAATCAAG |
| | Reverse | GGATAGTTGGGGAGCGACAC |
| *Adamts5* | Forward | GCATGCGCGCTACACTCTAA |
| | Reverse | TCGCGGTTGTAGACATGCAG |
